# Mechanical Fingerprints in Breast Cancer Research: A Multimodal Experimental Approach

**DOI:** 10.1101/2025.10.03.680332

**Authors:** Federica Banche-Niclot, Rosalia Ferraro, Valerio Di Palo, Paola de Paolis, Francesca Taraballi, Sergio Caserta

**Affiliations:** Center for Musculoskeletal Regeneration, Houston Methodist Academic Institute, Houston, TX, USA; Orthopedics and Sports Medicine, Houston Methodist Hospital, Houston, TX, USA; Dipartimento di Ingegneria Chimica, dei Materiali e della Produzione Industriale, Università degli Studi di Napoli “Federico II”, Napoli, ITA; CEINGE Advanced Biotechnologies Franco Salvatore, Naples, ITA

**Keywords:** Breast Cancer, Tumor Biomechanics, Viscoelastic Properties, 3D Spheroid Models, Tissue Stiffness, Translational Cancer Models

## Abstract

Breast cancer remains the leading cause of cancer-related mortality among women worldwide. Tumor biomechanics are not merely a symptom: they represent a functional signature with translational relevance in diagnostic, prognostic, and therapeutic resistance. Despite this, few experimental models are engineered to systematically investigate these physical properties across biological systems. Here, this study presents a multimodal biomechanical platform combining engineered 3D breast cancer spheroids with *ex vivo* tissue analysis to profiling and compare viscoelastic behavior or of healthy and tumoral environments.

Rheometry and compression testing revealed a consistent mechanical shift in tumor-derived samples marked by increased stiffness and force-dependent nonlinear behavior, mirroring the ECM remodeling typical of aggressive phenotypes. This increased rigidity may adversely affect chemotherapy effectiveness by hindering drug delivery and altering cellular mechanotransduction. These biomechanical fingerprints enable quantitative discrimination between healthy and cancerous tissues and can serve as a surrogate maker of malignancy. By supporting the development of mechanics-informed diagnostic tools, our platform offers a reproducible, clinically relevant framework to integrate biomechanical screening into translational breast cancer pipelines.

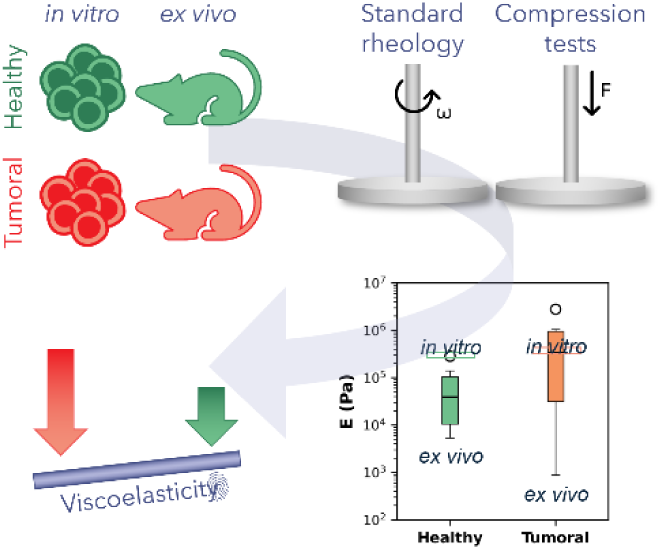

**Translational Impact Statement:** We propose a multimodal experimental approach that combines *in vitro* 3D breast-cancer models and *ex vivo* tissue analysis to measure and compare the viscoelastic properties of healthy and malignant breast tissues. By using mechanical behaviour as a fingerprint, this framework discriminates tumour tissue from its healthy counterpart. Revealing how tumour stiffness impacts drug delivery and therapy resistance, the approach provides a clinically relevant tool to inform diagnosis and optimise treatment strategies.

## 1. Introduction

Breast cancer is the most frequently diagnosed malignancy and a leading cause of cancer-related mortality among women worldwide [1]. While survival rates have improved for localized tumors thanks to early detection and localized treatments (such as surgery and targeted chemotherapy), metastatic breast cancer remains the leasing cause of death, accounting for over 90% of fatalities [2]. This highlights the critical and urgent need for models that capture molecular features along with the physical complexity of the disease.

In recent years, tumor biomechanics have emerged as a critical dimension of cancer biology. Mechanical alterations within the tumor microenvironment including extracellular matrix (ECM) stiffness, altered tissue viscoelasticity, and increased cellular deformability [3], are not passive byproducts of disease. Rather, increasing evidence shows that such biomechanical cues play an active role in reshaping cellular behavior and microenvironmental interactions [4], by actively influencing key hallmarks of cancer invasive and migrate, survive under stress, and respond to chemotherapeutic agents [5,6]. Tumor tissues often exhibit altered viscoelasticity and reduced compressibility, while cancer cells adapt through cytoskeletal remodeling and mechanical plasticity [7,8].

Despite this, mechanical phenotype remains underutilized in current diagnostic and therapeutic workflows and few platforms allow for controlled and quantitative exploration of these mechanical dynamics [9]. Conventional 2D cultures lack the spatial [10–13], heterogenicity [14,15] and mechanical context [16] of real tissues; while animal models offer limited tunability, scalability and translational accuracy [17]. 3D tumor spheroids (*i.e.*, multicellular compact aggregates) are gaining increased interest as a promising bridge, capable of recapturing the architecture, gradients, and cell-ECM interactions of native tumors as a scalable platform [18,19]. However, their biomechanical characterization remains challenging and underexplored. *Ex vivo* experiments, which involves tissues or organs from living organisms, can come into play allowing also the maintenance of tissue natural architectures and cell-to-cell interactions, minimizing alteration to the living environment and thus offering a more realistic approximation of *in vivo* conditions [20]. Nevertheless, a clinically relevant *in vitro* model must be quantitatively validated against real tissue mechanics using reliable and physiological meaningful techniques.

Traditional techniques such as atomic force microscopy (AFM) [21–23], indentation [24], micro tweezers [25], and compression assay [26,27] have contributed valuable insights into cellular elasticity, yet they often oversimplify the inherently viscoelastic nature of biological systems. These methods are sensitive to tool geometry and substrate properties, resulting in high variability and limited comparability especially when transitioning from isolated cells to tissue-level constructs. In contrast, rheology provides a robust and scalable method to quantify viscoelastic demeanor in both soft solids and complex biological matrices. By measuring elastic (G′) and viscous (G″) moduli, rheology identify both energy storage and dissipation which are key parameters for understanding how tumors deform, resist, or transmit mechanical stress. These moduli can be translated into conventional mechanical metrics (*e.g.*, Young’s modulus) under defined assumptions, allowing comparisons across biological models [28]. Given the deformability, heterogeneity, and flow-like behavior of cancerous tissues, rheological analysis offers a physiologically relevant and reproducible approach for characterizing tumor mechanics at experimental scales.

In this study, we present a multimodal experimental strategy that couple *in vitro* spheroids with *ex vivo* tissue analysis to quantify the viscoelastic profiles of healthy and cancerous breast environments. Using rotational rheometry and compression testing, we mapped the viscoelastic landscape of breast cancer and healthy controls.

Our approach aims to establish a reproducible mechanical benchmark that distinguishes mechanical alterations associated with tumor pathology, laying the foundation for incorporating biomechanics into translational breast cancer research with potential application in both diagnostic development and therapy optimization.

## 2. Materials and Methods

### Cell Culture and Spheroids Formation

The biological component of this study relied on two distinct cell lines, organized into spheroids to better replicate the three-dimensional tumour microenvironment. The first model system consisted of MDA-MB-231 human breast adenocarcinoma cells (official designation: MDA-MB-231, HTB-26; Homo sapiens, female; pleural effusion from a patient with metastatic breast adenocarcinoma; RRID:CVCL_0062), obtained from the American Type Culture Collection (ATCC, Manassas, VA, USA) in March 2022. ATCC authenticates this cell line by short tandem repeat (STR) profiling, and MDA-MB-231 is not listed as misidentified or cross-contaminated in the International Cell Line Authentication Committee (ICLAC) register.

As a control, primary human mesenchymal stem cells (FP-MSCs) were isolated from infrapatellar fat pad tissue (male donor, 24 years old, height 188 cm, weight 72.6 kg, BMI 20.5) under institutionally approved protocols (Houston Methodist Hospital IRB Pro00015718) in November 2022). FP-MSCs were characterized by the institutional biorepository using standard MSC immunophenotyping (CD73+, CD90+, CD105+, CD45−) prior to experimental use. Because FP-MSCs are primary cells rather than commercial or immortalized lines, a Research Resource Identifier (RRID) is not applicable.

Mycoplasma testing was not performed on either MDA-MB-231 or FP-MSC during this study. While this represents a potential limitation, no morphological abnormalities or altered growth kinetics were observed, and the reproducibility of biomechanical results across replicates supports that contamination did not influence the experimental outcomes.

The MDA-MB-231 were cultured in Dulbecco’s Modified Eagle’s Medium (DMEM) supplemented with 20% (v/v) Fetal Bovine Serum (FBS) and 1% (v/v) antibiotics (50 U/mL penicillin and 50 mg/mL streptomycin). Cells were used at passages 3–9. On the other hand, FP-MSC was obtained from the orthopedic biorepository at Houston Methodist Hospital, which were previously obtained by the Department of Orthopedics and Sports Medicine Department (IRB Pro00015718). Those were cultured in alpha Minimum Essential Medium (α-MEM) supplemented with 20% (v/v) FBS and 1% (v/v) antibiotics. Cells were used at passages 1–6. Both cell lines were cultured under sterile standard conditions (37°C, 5% CO_2_). Transiting from 2D to 3D cultures, cellular spheroids were prepared using the liquid overlay technique [29], in which a non-adhesive concave surface was employed to promote cell-cell aggregation within the meniscus. Specifically, the nonstick polymer film on the wells was obtained by coating U-bottom shaped 96-well plate (Corning TM Costar TM 96-Well, Cell Culture -Treated, U-Shaped Bottom Microplate, Thermo Fisher Scientific) with 0.1% polyHEMA in 96% ethanol. Cells were then seeded at a concentration of 2·10^3^ cells/well. FP-MSC spheroids were cultured in the medium described above, whereas MDA-MB-231-GFP ones were maintained in an optimized medium (DMEM with 1% antibiotics, 2% bovine serum albumin (BSA), 0.05% insulin, and 1% Matrigel). Additionally, the cancer multiwell plate was placed on a rotating plate to enhance the cells aggregation. Both spheroids were then incubated under typical cell culture conditions to obtain compact spheroids of adequate size: 1 day for FP-MSCs and 3 days for MDA-MB-231-GFPs.

### Cell Viability Assessment

Two complementary assays were employed to evaluate spheroids viability over time. Specifically, at designated time points (24, 48, and 72 hours), the ATP-related metabolic activity of spheroids was quantified through CellTiter-Glo® 3D Assay (Promega Corporation, Madison, WI, USA) following manufacturer’s instruction. Alongside, live and dead cells were determined via Live/Dead fluorescent staining by using two fluorescence probes that simultaneously color intracellular esterase activity and plasma membrane integrity, recognized parameters of cell viability. For this purpose, cell-permeant calcein AM (1 μM, Thermo Fisher Scientific) and ethidium homodimer-1 (EthD-1, 2 μM, Thermo Fisher Scientific) were used to stain live cells (green fluorescence) and dead cells (red fluorescence), respectively. Samples were incubated with the staining solution for 15 minutes before imaging. Experiments were performed in triplicate.

### Collagenous supporting hydrogel preparation

Collagen-based support with suitable stiffness was developed to stabilize spheroids during rheological analysis and thus preventing uncontrolled movement and ensuring an adequate gap for accurate measurements. For this aim, a purified methacrylated Type I bovine collagen (PhotoCol® Methacrylated Collagen, CMA, Advanced BioMatrix) solution was prepared at a concentration of 3 mg/mL following the manufacturer’s instructions. To produce a uniform collagen matrix for supporting spheroids during rheological characterization, 2 mL of collagen solution was poured into a circular silicon mold of 25 mm diameter and a thickness of 2.5mm, matching the size and shape of the measurement tool. The samples were then incubated at 37°C with 5% CO_2_ for one hour to ensure physical crosslinking of collagen.

### Assembly of Collagen-Spheroid Constructs

Cellular spheroids were placed on the crosslinked collagen disk before rheological characterization. The number of spheroids was selected to achieve approximately 1% surface coverage (S%) of the supporting disk, defined as n · (A_s_⁄A_gel_) · 100 = n · (d_s_⁄d_gel_) · 100. Here, n is the number of spheroids, d_s_ the mean spheroids diameter for each cell lines, and d_gel_ the diameter of the collagenous hydrogel (25 mm). The average diameter of the MDA-MB-231-GFP spheroids was calculated to 580.00 ± 30.55 μm, while 256.66 ± 14.53 μm for FP-MSC spheroids (**Figure S1**). Based on these values, 15 spheroids were required for MDA-MB-231-GFP cells, whereas 85 spheroids were needed for FP-MSC cells. Therefore, spheroids were individually collected from the 96-well plates and dropwise on the hydrogel’s surface, ensuring a uniform distribution. Due to the high number of FP-MSC spheroids required, they were arranged in groups of approximately 10 spheroids each along the matrix’s surface.

### Tissues manipulation

To evaluate the physiological relevance of the developed *in vitro* spheroid-based 3D models, the biomechanical properties of murine adipose tissue and tumor masses were examined and compared to those of spheroids. Given that animal models remain the primary established system for studying tumor mechanics, this comparison aimed to assess whether the *in vitro* model reliably recapitulates key biomechanical features observed *in vivo*. To this aim, mammary gland fat tissue from mice was analyzed as the *ex vivo* equivalent of FP-MSC spheroids, while tumors generated by MDA-MB-231 cell injections into mice served as a reference for MDA-MB-231-GFP spheroids. The sample tissues were thawed using 1x Phosphate-Buffered Saline (PBS) and transferred to a Petri dish, where they were left to rehydrate for a minimum of 5 minutes to ensure optimal restoration of their physiological properties. Prior to testing, the tissues were removed from the PBS, and if necessary, standardized to a 5 mm diameter using a biopsy punch to ensure consistency in sample size and facilitate reproducible mechanical measurement.

### Rheological setup

Oscillatory measurements of both *in vitro* and *ex vivo* systems were carried out by using a stress-controlled rheometer (Anton Paar Physica MCR 301 Instrument) equipped with a smooth parallel plate measuring system with a diameter of 25 mm (measuring plate PP25-25MM, Anton Paar). For guaranteeing spheroids viability by mimicking physiological condition, the temperature of the rheometer’s lower plate was kept at 37°C using a Peltier system. Additionally, to prevent water evaporation and to ensure a uniform temperature around the sample, the rheometer plate was enclosed within a chamber to create a controlled environment. The viscoelastic properties of systems were assessed by performing strain (γ) sweep tests, with amplitudes ranging from 0.05% to 1% at a constant angular frequency (ω) of 10 rad/s, to identify the linear viscoelastic range. Elastic and viscous moduli (G’ and G’’, respectively) were measured by varying ω within the range 0.01 – 100 rad/s.

### Compression setup

Compression tests were performed on *ex vivo* tissues by using a mechanical testing system (Univert Cell Scale biomaterials testing) equipped with a 1 N load cell. Measurements were performed on both tumor and non-tumor tissues, which were standardized to a cylindrical shape with a diameter of 5 mm. The mechanical setup was designed to assess the viscoelastic behavior of the tissues, capturing its response to deformation and short-term recovery under controlled mechanical stress. For achieve this, a single compression cycle of 3 minutes, followed by a 5-second recovery period, was applied with a stretch magnitude of 50%, ensuring a standardized assessment of the tissue’s mechanical behavior. Mechanical properties of the samples were expressed in terms of Young’s modulus (E), determined from the slope of the linear correlation between stress (σ) and strain (ε) [30,31].

### Statistical analysis

Cellular assessments were performed in triplicates and results were analyzed through two-way ANOVA method (unpaired, two-tailed Welch’s t-test, n=4). Rheological measurements were performed at least twice for each system, while compression tests were repeated eight times for tissues type. Data are presented as mean ± standard error of the mean (SEM), where SEM is defined as the standard deviation of the population normalized by the square root of the sample size.

## 3. Results

### Spheroids viability

To assess the most appropriate spheroids formation protocol, viability of both FP-MSC and MDA-MB-231-GFP spheroids was evaluated using CellTiter-Glo® 3D Assay over a 72-hour period. In particular, optical imaging confirmed the morphologies in 2D monolayer and spheroids set-up (**Figure 1a**) while **Figure 1b** reported the metabolic activity of the 2D control compared with spheroids, healthy cells in the upper histogram and the cancer cells in the lower one. From both cell lines, the spheroids metabolic activity was significantly lower compared to 2D cultures, particularly for FP-MSC condition. This reduced viability in spheroids is likely a result of the physical constraints of their architecture and the formation of a necrotic core, which deprives the central cells of nutrients and oxygen. In contrast, 2D controls exhibited a continuous increase over time, reflecting the proliferation of cells in the bidimensional configuration. Conversely, spheroids are characterized by a non-proliferative nature and their cells progressively consolidating around the core, as demonstrated by the moderate downward trend over time, particularly for FP-MSC ones. Despite these differences, spheroids result alive after three days of culturing and the observed trend can be ascribed to the expansion of the necrotic core, an intrinsic feature of this type of 3D construct [17], as demonstrated by Live/Dead staining (**Figure 1c**).

**Figure 1.**
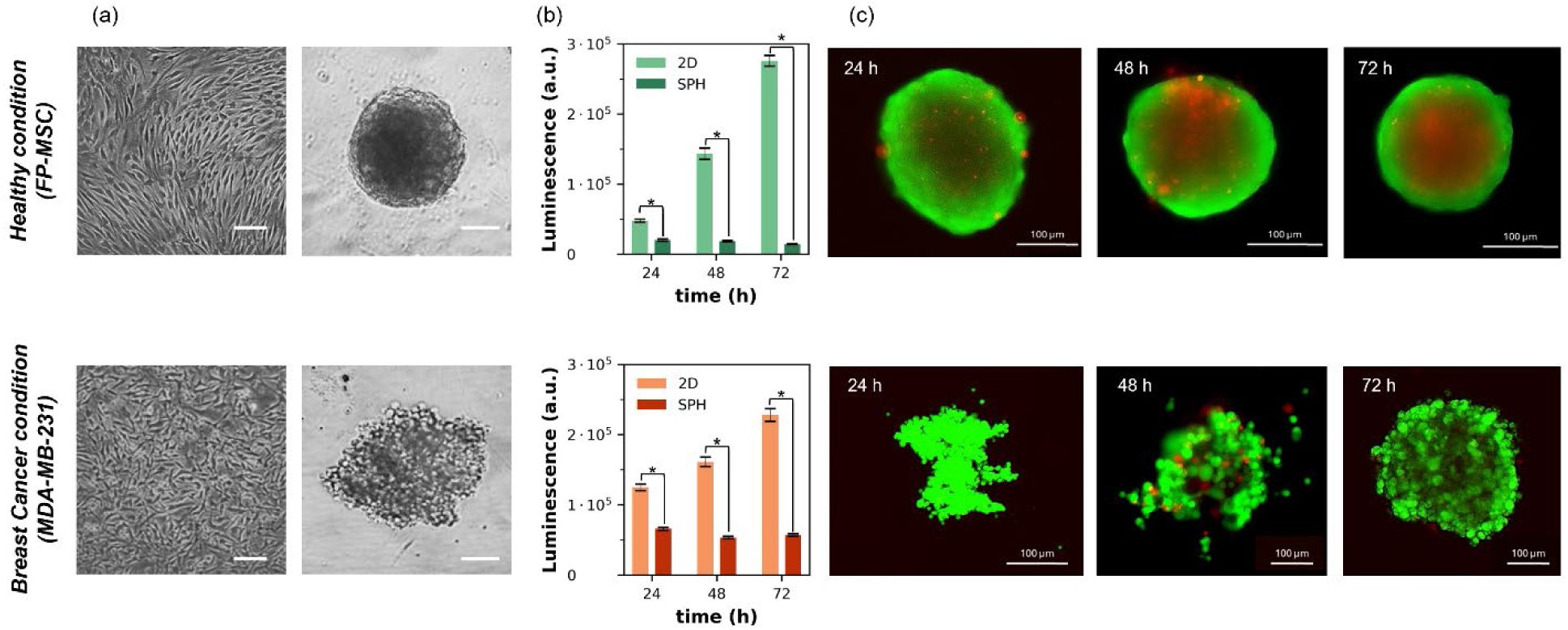
Spheroids morphology and viability investigations: **a**) Optical images of 2D monolayer (left) and spheroids (right) for FP-MSC (above) and MDA-MB-231-GFP (below). Scale bar: 100 μm. **b**) Live/Dead images of FP-MSC spheroids (above) and MDA-MB-231-GFP (below) spheroids at different timepoints (24, 48 and 72 hours). **c**) Cells viability assessment on both the 2D control and the spheroids configurations at different timepoints (24, 48 and 72 hours). Statistical significance of the results: *p<0.05.

Considering the FP-MSC spheroids images (**Figure 1c**, top row), they initially exhibited a compact structure with mixed fluorescence, with green indicating live cells and red indicating dead cells. However, as time progressed, a clear necrotic core began to form, particularly evident after 72 hours. This was reflected by the increased red fluorescence, indicating a higher number of non-viable cells. Given the higher metabolic activity and the absence of a necrotic core, FP-MSC spheroids were tested after 24 hours to ensure their viability.

Different behaviors were observed for MDA-MB-231-GFP spheroids. Initially, the cells are unable to aggregate into a well-defined 3D structure at 24 and 48 hours. A nearly spherical shape morphology was observed only at 72 hours. Additionally, the development of a necrotic core moderately starts to develop at the last timepoint. Despite the onset of necrosis, the metabolic activity remained nearly constant over the 72-hour experimental period was not significantly influenced by the necrotic core. Therefore, due to the sustained viability at 72 hours and the absence of a fully developed necrotic core, the mechanical properties of MDA-MB-231 spheroids were assessed at this timepoint to ensure proper spheroid formation.

### Mechanical characterization of in vitro 3D models

Rheological analyses were performed on the *in vitro* 3D models, using collagen-based hydrogel as supporting material (**Figure 2**). FP-MSC (NT-S) and MDA-MB-231-GFP (T-S) spheroids were embedded in the collagen-based hydrogel (COL) (**Figure 2a**). Bright field/fluorescence imaging confirmed a uniform spheroid distribution throughout the supporting material (**Figure 2b**). Samples were then mounted in a plate–plate rheometer for amplitude oscillatory testing (**Figure 2c**). To ensure a comprehensive evaluation, the viscoelastic properties of the collagenous hydrogel (COL) were also characterized independently as a reference. **Figure 2d** presents the rheological outcomes for non-tumoral (NT-S, composed of FP-MSC) and tumoral (T-S, composed of MDA-MB-231-GFP) spheroids, highlighting key differences in their viscoelastic behavior in terms of G’ and G’’ (represented by upward and downward triangles, respectively) reported as function of ω. As shown, the viscoelasticity properties of pure collagen (data reported in blue) are approximately three orders of magnitude lower than those of the system containing spheroids (COL+NT-S in green and COL+T-S in orange, respectively). At ω∼1 rad/s, G’ for the supporting hydrogel alone is approximately 78 Pa, whereas for the *in vitro* 3D systems, G’ reaches 9.70±1.83·10^4^ Pa for COL+NT-S and 2.45±1.13·10^5^ Pa for COL+T-S. This confirms that collagen stiffness does not influence the mechanical properties of the spheroids, making it a suitable support material that mimics the soft ECM environment around the cells. Interestingly, the biomaterial alone exhibits a crossover point (G’ = G’’) at ω∼45 rad/s, indicating a transition in its mechanical behavior from solid-like to liquid-like. Contrarily, no crossover is observed in the samples containing spheroids, suggesting a persistently dominant elastic response throughout the tested frequency range and reinforcing the solid-like nature of spheroids. In addition, the incorporation of spheroids enhanced the structural integrity and elastic nature of the construct. This aligns with the behavior of native tissue, which exhibits a predominantly elastic response due to its ECM composition and cell-cell interactions [32]. Thus, these results highlight the importance of cellular components in maintaining the elastic nature of tissue and confirm that the spheroid-laden hydrogel more effectively mimics the biomechanical properties, providing a more physiologically relevant *in vitro* model for studying tissue mechanics. Focusing on the *in vitro* 3D models, tumoral spheroids exhibited a higher G’ compared to healthy ones (approximately 2.45·10^5^ Pa and 9.70·10^4^ Pa, respectively, at ω∼1 rad/s). Particularly, the G’ values of these spheroids were consistent across collagen concentrations, further suggesting that the mechanical properties of FP-MSC spheroids are independent of the supporting matrix stiffness within this range [33,34]. In stark contrast, tumoral MDA-MB-231-GFP spheroids exhibited significantly higher stiffness, with a G’ value 2.5 times greater than that of FP-MSC spheroids, which is consistent with the widely observed phenomenon of tumor tissues being mechanically stiffer than healthy tissues. This increase in stiffness was accompanied by a higher value of tan δ, which is indicative of a more solid-like behavior in tumor spheroids compared to their healthy counterparts.

**Figure 2.**
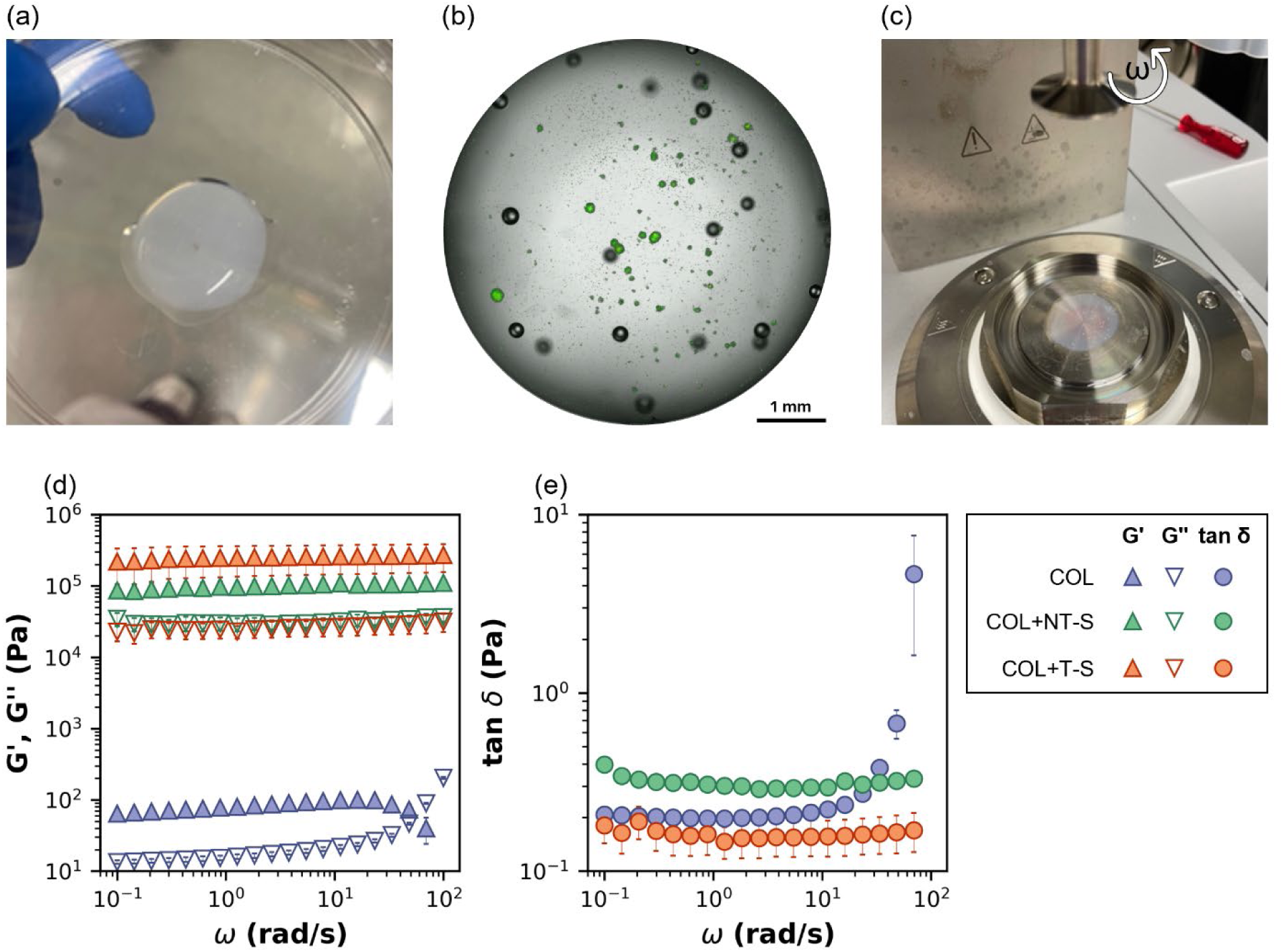
Sample preparation and viscoelastic properties of FP-MSC (NT-S) and MDA-MB-231-GFP (T-S) spheroids using collagen-based hydrogel (COL) as supporting material. **a**) Representative image of the COL hydrogel disk after gelation, embedding cellular spheroids prior to rheological measurements. **b**) Representative bright-field/fluorescence image showing GFP-positive T-S distribution within COL. Scale bar: 1 mm. **c**) Rheometry setup: COL hydrogel with embedded spheroids loaded in a plate–plate geometry for oscillatory testing. **d**) Viscous (G’, Pa) and elastic (G’’, Pa) moduli as function of angular frequency (ω, rad/s) for collagenous hydrogel without and with spheroids non-tumoral or tumoral (blue for bare supporting hydrogel, green with non-tumoral spheroids (COL+NT-S) and orange with tumoral spheroids (COL+T-S)), represented by upward triangles and downward triangles, respectively. **e**) Tan δ as function of angular frequency (ω, rad/s) for collagen-based hydrogel without and with spheroids, represented by circles.

Indeed, tan δ (*i.e.*, the ratio between G’’ and G’) was analyzed for providing additional insights into the viscoelastic behavior of the system and a more comprehensive characterization of the biomechanical fidelity of the *in vitro* model. **Figure 2e** shows tan δ as function of ω, where biomaterial alone is represented in blue circle, COL+NT-S in green and COL+T-S in orange. For the supporting material, tan δ remains nearly constant at approximately 0.20 for ω<10^1^ rad/s. However, for ω>10^1^ rad/s, tan δ starts increasing, reaching a final value of approximately 4.64, indicating the transition from solid-like to liquid-like behavior. In contrast, for the COL+NT-S, tan δ remains constant at approximately 0.30 across the entire frequency range and, similarly, for the COL+T-S, it stays nearly constant at around 0.15. Consequently, tumoral spheroids exhibited a more pronounced solid-like behavior compared to their non-tumoral counterparts.

To gain deeper insight into the biomechanics of the *in vitro* 3D models, E was evaluated for all tested systems. The E values were obtained by considering the G’ value at ω∼1 rad/s and applying the relationship E=2G’(1+ν), where ν is the Poisson ratio, set to 0.5. In particular, for incompressible materials such as fat, ν is typically assumed to be 0.5. Specifically, for fat pad tissue, ν≈0.475, while for breast fat, it is reported around 0.49 [35]. In contrast, cancerous tissues generally exhibit lower Poisson’s ratios and ν ranges between 0.2 and 0.45 for most breast cancer tissues, depending on the subtype. The most common forms of breast cancer typically show values around 0.3–0.35, whereas triple negative breast cancer (TNBC) is considered nearly incompressible, with ν≈0.5 [36,37]. As a result, the calculated E values are 2.34±0.16·10^2^ Pa for supporting hydrogel alone, 2.91±0.55·10^5^ Pa for COL+NT-S and 4±0.44·10^5^ Pa for COL+T-S. These values confirm the significant difference in stiffness (three orders of magnitude larger) when spheroids are integrated, with tumoral spheroids exhibiting a much stiffer behavior compared to their healthy counterparts. Such mechanical alterations are typically associated with the ability of cancer cells, especially those in more aggressive phenotypes, to remodel the ECM, including increased collagen crosslinking and altered ECM composition, which collectively contribute to tumor rigidity increasing their contractile forces as well as promote their migration and invasive capabilities [38–40]. These enhanced mechanical properties of MDA-MB-231-GFP cancerous spheroids reflects this remodeling valuable insights into the altered biomechanical behavior of cancer cells and their role in metastasis [41].

### Ex vivo tissues biomechanics

To benchmark against *in vitro* 3D models, *ex vivo* mammary gland fat pad (healthy) and MDA-MB-231 tumor-bearing tissues were excised (**Figure 3a**) and standardized with 5-mm punches (**Figure 3b–c**). To assess the reliability of the *in vitro* models, the biomechanical properties of *ex vivo* healthy and tumoral tissues were investigated through rheological analyses and compression analyses. **Figure 3d** presents the rheological outcomes for healthy and tumoral tissues, highlighting key differences in their viscoelastic behavior in terms of G’ and G’’ (represented by upward and downward triangles, respectively) as function of ω. As observed in the *in vitro* 3D models, the tumoral tissues (orange symbols) are stiffer than the healthy counterpart (green symbols), with G’ values of 2.38±1.04·10^5^ Pa and 2.45±0.93·10^4^ Pa at ω∼1 rad/s, respectively. Both systems exhibit solid-like behavior, with G’ consistently higher than G’’, specifically tumoral tissues are approximately 10 times stiffer than their healthy counterparts.

**Figure 3.**
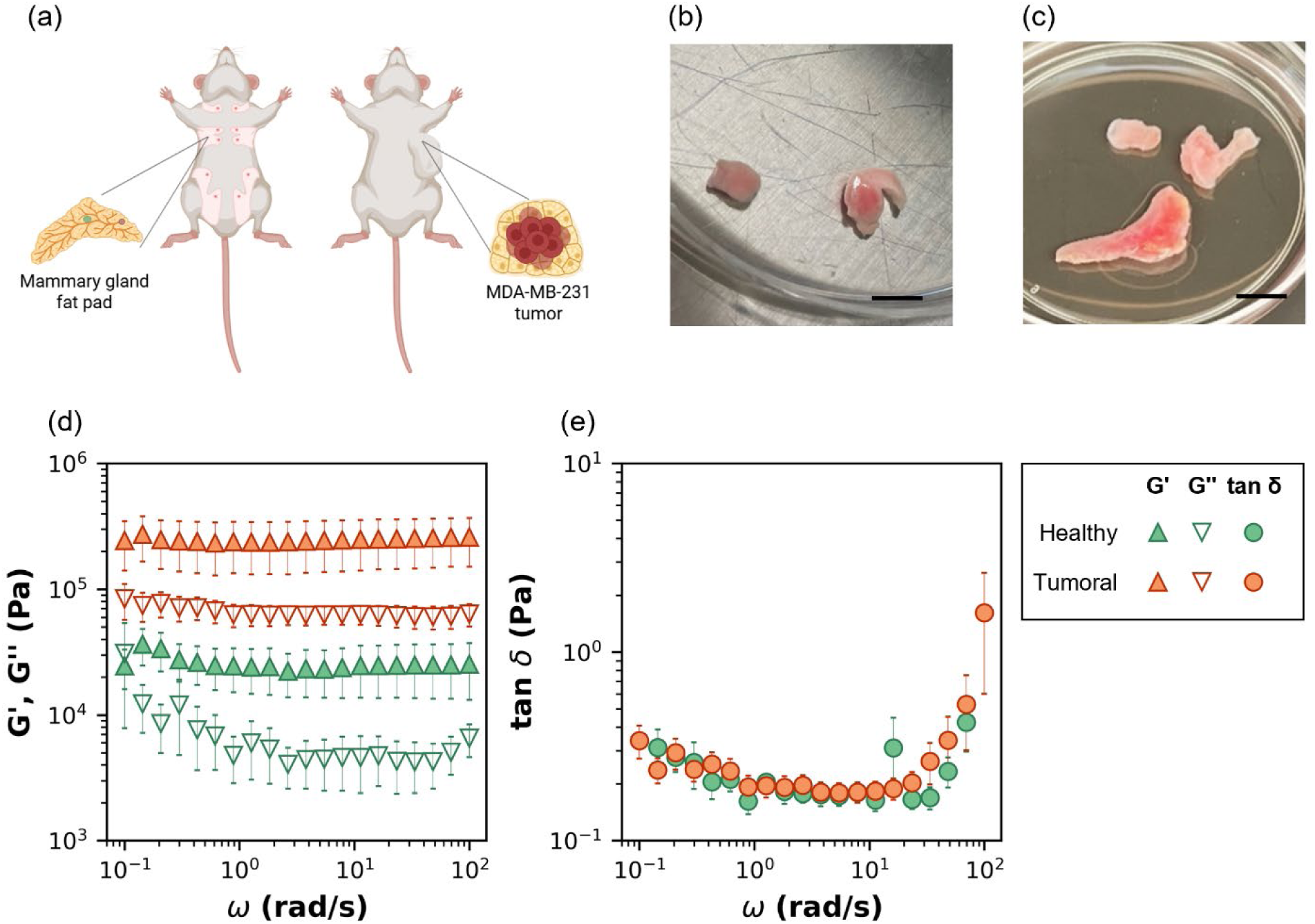
Extraction, sampling and viscoelastic properties of healthy and tumoral tissues. **a**) Schematic of tissues extraction: mammary gland fat pad used as healthy control (left) and MDA-MB-231 tumor-bearing tissue used as tumoral condition (right). **b**) Representative healthy explants after resection and cored with a 5-mm biopsy punch. Scale bar: 5 mm **c**) Representative tumoral explants excised from mice bearing MDA-MB-231 lesion after resection and cored with a 5-mm biopsy punch. Scale bar: 5 mm. **d**) Viscous (G’, Pa) and elastic (G’’, Pa) moduli as function of angular frequency (ω, rad/s) for healthy and tumoral tissues (green and red symbols, respectively), represented by upward triangles and downward triangles, respectively. **e**) Tan δ as function of angular frequency (ω, rad/s) for healthy and tumoral tissues (green and red circles, respectively).

Figure 3e further supports this finding by displaying the evolution of tan δ as a function of ω. Unlike the *in vitro* model, where tan δ for the healthy system was higher than for the tumoral system, here the values are quite similar for tissues regardless of pathology (0.16 *vs.* 0.19 for healthy and tumoral tissue, respectively) for ω<10^1^ rad/s. At higher frequencies, tan δ increases up to approximately 1.61.

Additionally, E was evaluated for all tested systems by considering the G’ value at ω∼1 rad/s and assuming ν ≈ 0.49 for breast fat [35] and 0.37 for tumoral tissue [36,37]. Healthy tissues exhibit an E value of 7.30±2.77·10^4^ Pa, whereas tumoral tissues display a value one order of magnitude higher (6.43±2.97·10^5^ Pa). This trend closely aligns with the developed *in vitro* 3D model results, further confirming the biomechanical relevance of the *in vitro* approach. Interestingly, the variability in the results is more pronounced in the *ex vivo* samples, particularly for tumoral tissues, and this aspect will be further clarified in the following section.

To validate the biomechanical findings on *ex vivo* samples, E values were also obtained using a biological compressor for both non-tumoral and tumoral tissues. Specifically, the Hertz model was employed to calculate E from the slope of the stress–strain (σ–ε) curve, considering only strains below 10% to ensure the analysis remained within the linear elastic regime [42]. As shown in Figure 4a, tumoral tissues (shown in orange) exhibit an E value of 5.22±0.62·10^4^ Pa, while their healthy counterpart (in green) shows a lower value of 3.68±0.38·10^4^ Pa. This indicates that tumor tissue is approximately 1.42 times stiffer, consistent with the rheological outcomes observed for both the optimized *in vitro* 3D models and the *ex vivo* samples. Additionally, the variability observed in the compression measurements is lower than that in standard rheology (Figure 4b), likely due to the smaller variation in the applied normal force (F_N_), while maintaining a constant cross-sectional area.

**Figure 4.**
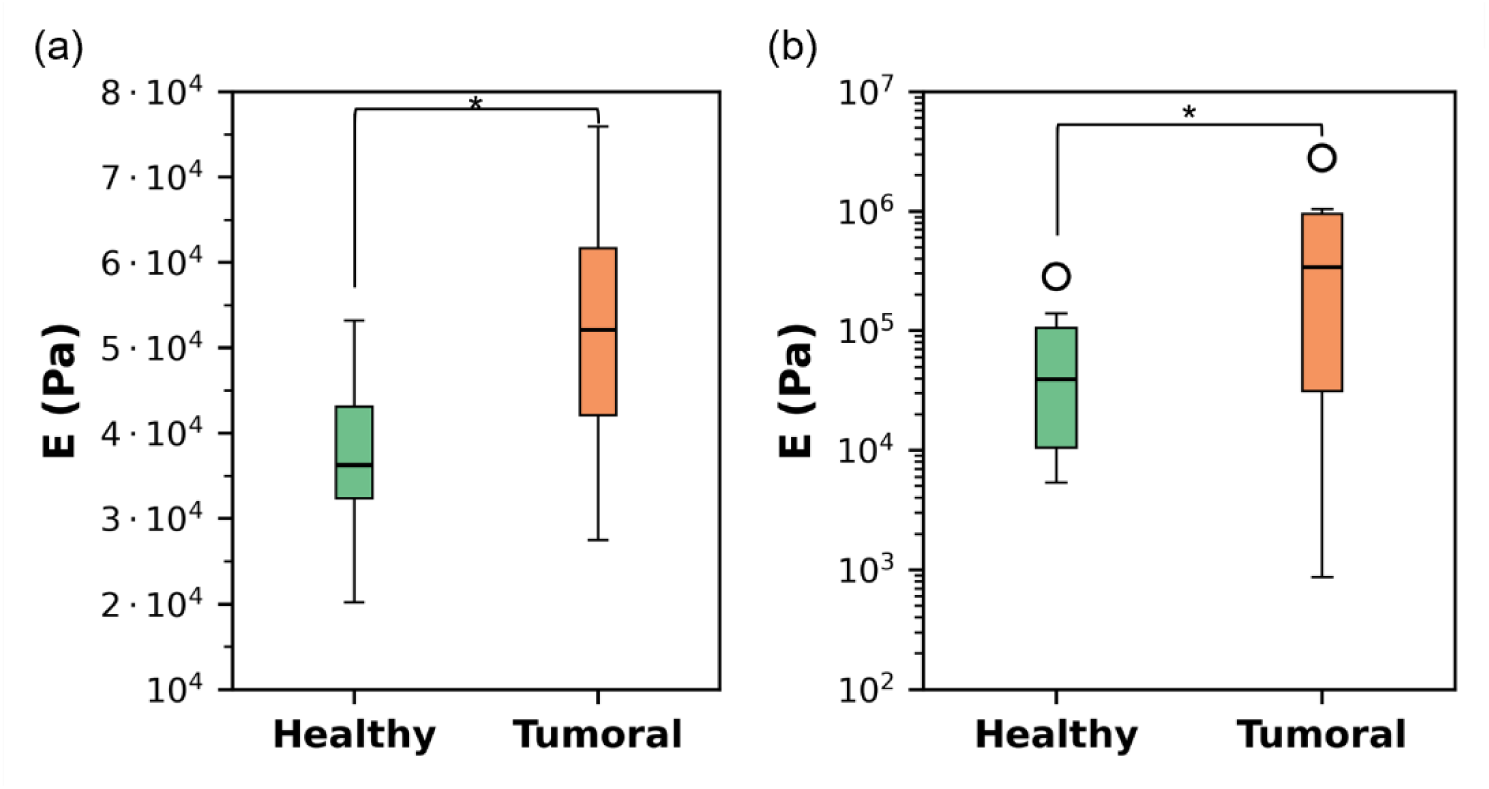
Mechanical evaluation of ex vivo tissues. Comparison of the Young’s modulus (E) for non-tumoral and tumoral tissues (green and orange, respectively) in (**a**) compression tests and (**b**) standard rheology. Statistical significance of the results: *p<0.05.

Notably, depending on the pathological status, biological samples may exhibit either force-softening (healthy tissues) or force-stiffening (tumoral tissue) behaviors. Such non-linear responses can introduce artifacts into the characterization process, compromising a direct correlation between the viscoelastic properties of ECM and critical cellular behaviors, such as migration, proliferation, or differentiation [43,44]. However, when accounting for the normal force applied, a very good agreement between mechanical outcomes can be achieved. In fact, the discrepancy in the absolute values of E between rheological and compression-based measurements (*i.e.*, 7.30±2.77·10^4^ Pa *vs* 3.68±0.38·10^4^ Pa for healthy tissues, respectively; and 6.43±2.97·10^5^ Pa *vs* 5.22±0.62·10^4^ for tumoral tissues, respectively) can be attributed to differences in the applied normal force. During standard rheological testing, samples were intentionally compressed to ensure full contact between the biological tissue and the rheometer plates (*i.e.*, d_tissues_∼d_tool_) thereby minimizing measurement artifacts [43,44]. Due to sample variability, different normal forces were applied along the tissue surface, resulting in variations in the measurement gap, typically ranging from 0.1 to 0.7 mm for healthy tissues and from 0.3 to 1.0 mm for tumoral tissues. Figure 5 display the elastic (G′, solid upward triangles) and viscous (G″, open downward triangles) moduli for both healthy and tumoral tissues (Figure 5a and **5c**, respectively), plotted as a function of the applied F_N_ while keeping the cross-sectional area constant and approximated by the tool area (A = πR^2^ ∼ 491 mm^2^). In all conditions, G′ consistently exceeds G″, confirming the predominantly elastic nature of the tissues. Interestingly, healthy tissues (Figure 5a) exhibit a force-softening response, with moduli decreasing by at least one order of magnitude as the F_N_ increases. In contrast, tumoral tissues (Figure 5) show a force-stiffening behaviour, whereby higher force leads to increased moduli.

**Figure 5.**
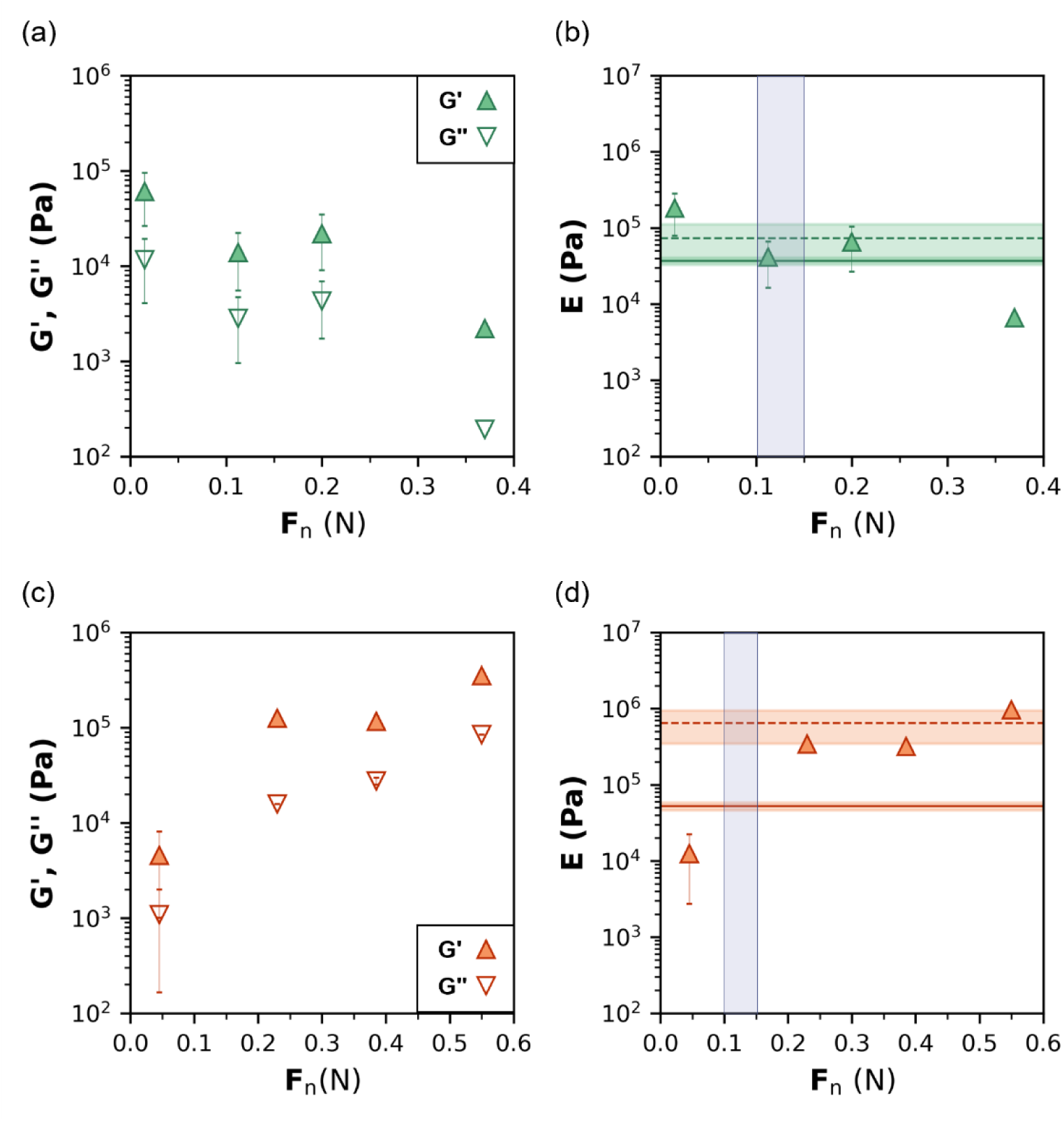
Viscoelastic properties of healthy and tumoral tissues as a function of applied normal force (F_N_). **a,c**) Elastic (G’, Pa) and viscous (G’’, Pa) moduli plotted against applied stress (σ, Pa) for healthy (green triangles) and tumoral (red triangles) tissues, represented by upward triangles and downward triangles, respectively. **b,d**) Comparison of the Young’s modulus (E) for healthy (green) and tumoral (red) tissues, estimated using the relation E = 2G’(1 + ν), with G’ evaluated at ω ≈ 1 rad/s. In both panels, the mean E values obtained from standard rheological measurements (dotted line) and from biological compression tests (solid line) are indicated.

To better compare the results of standard rheology with those from the compression assay, **Figure 5b** and **5d** show the corresponding E, calculated by assuming the previously discussed *ν* (0.49 and 0.37 for healthy and tumoral tissues, respectively), plotted as a function of F_N_. In both plots, the average E value obtained *via* standard rheology is marked by dotted line (7.30·10^4^ Pa and 6.43·10^5^ Pa for healthy and tumoral tissues, respectively). During compression testing, the F_N_ reached for strain ε < 10% was approximately 0.1 – 0.15 N. Within this F_N_ range, the predicted E values from rheology closely match those measured via compression (3.68·10^4^ Pa and 5.22·10^4^ Pa for healthy and tumoral tissues, respectively), indicated by solid lines. This agreement allows for a more reliable comparison of tissue elasticity across the two methods. Notably, the discrepancy observed between rheology and compression-based values, more pronounced in tumoral tissues, is effectively resolved when actual F_N_ is accounted for, demonstrating strong consistency between the datasets. This highlights the importance of comparing mechanical properties within similar normal force regimes to ensure consistency between characterization methods.

## 4. Discussion

Understanding and quantifying tumor biomechanics is gaining momentum as a critical dimension of cancer research, with growing evidence that mechanical alterations within the tumor microenvironment actively influence tumor progression, metastasis, and resistance to therapy. Despite this recognized importance, tools capable of capturing and comparing tumor mechanical properties across biological systems remain limited. In this context, the platform presented in this study offers a technologically robust and biologically relevant solution, bridging 3D *in vitro* spheroids with *ex vivo* tissue analysis through a unified rheological and compression-based approach.

Our findings confirm that breast cancer spheroids derived from MDA-MB-231 cells exhibit significantly higher stiffness and nonlinear force-dependent behavior compared to their non-tumoral counterparts, reflecting ECM remodeling associated with aggressive phenotypes. Moreover, the choice of a collagen-based hydrogel ensured that the supporting matrix did not mask spheroid mechanics while maintaining a biomimetic environment. The difference between condition was preserved across *in vitro* and *ex vivo* models, with tumor tissues showing elevated storage modulus (G′) and distinct nonlinear behavior under load, both signatures of ECM remodeling and increased cellular contractility. The consistency between *in vitro* and *ex vivo* measurements across force regimes supports the mechanical fidelity and transferability of our *in vitro* system and validates its potential as a preclinical platform for studying or testing mechano-sensitive tumors and stiffness-adapted therapies. While further validation is required, this work lays the groundwork for integrating mechanical parameters into diagnostic workflows and therapeutic planning.

Importantly, this study highlights the complementary nature of rheological and compression-based assessments. Technologically, the choice of rotational rheometry as a central characterization tool marks a critical advantage. While compression testing remains standard in tissue mechanics, they often oversimplify tissue mechanics by focusing solely on elastic properties [45], typically through the application of the Hertz model. In contrast, rheology provides a richer, frequency-dependent profile of viscoelastic behavior, critical for capturing the dynamic nature of tumor response. The combination of rheological and mechanical compression techniques highlights the strengths and complementary nature of these techniques, with rheology providing a more comprehensive understanding of material behavior at both the cellular and tissue levels. The agreement between these methods within specific normal force windows emphasizes the need for careful comparison of mechanical properties under consistent conditions. Our data shows that force-sensitive shifts in stiffness can significantly affect measurements and should be standardized or accounted for in future biomechanical profiling.

From a translational standpoint, the observed mechanical differences may have direct implications for therapy design and diagnostics. Increased tumor stiffness has been shown to hinder drug diffusion, impair immune cell infiltration, and activate mechanotransduction pathways that promote chemoresistance [5,6]. By reproducing this stiffness landscape *in vitro* and aligning it with *ex vivo* profiles, our system enables a more predictive evaluation of mechanically responsive therapies. Moreover, the mechanical fingerprinting of tumor models could support the development of diagnostic criteria based on physical properties, especially when integrated with imaging techniques or functional assays.

Critically, the reproducibility and scalability of our platform make it suitable not only for academic investigation but also for industrial applications in drug screening and biomaterials testing. The use of standardized hydrogel scaffolds, controlled spheroid assembly, and quantitative mechanical output positions this approach as a translatable solution for early-stage therapeutic validation or personalized medicine pipelines. While further work is needed to expand the system to other tumor subtypes and integrate dynamic environmental conditions (e.g., perfusion, immune components), the current framework offers a solid starting point for embedding biomechanical profiling into preclinical workflows.

In summary, the platform presented here combines engineered biological models with precision mechanical analysis, offering a scalable and reproducible tool for translational breast cancer research. By bridging tissue-level biomechanics and experimental oncology, this approach opens new directions for mechanics-informed diagnostics, therapy design, and patient-specific modeling.

## 5. Conclusion

This study presents a robust, scalable platform for biomechanical profiling of breast cancer using a multimodal strategy that bridges 3D *in vitro* spheroids and *ex vivo* tissue mechanics. Our approach bridges scalable *in vitro* models and complex tissue environments, providing a quantitative insight into viscoelastic tumor behavior, an often-overlooked dimension with growing clinical relevance.

The ability to reproduce and compare mechanical signatures across biological systems supports applications in drug screening and, mechanics-informed therapy design, and physical biomarker discovery, aiming to improve drug delivery and treatment efficacy in cancer patients. Furthermore, given the importance of ECM stiffness in cancer biology as a predictor of therapeutic resistance and disease progression, our platform offers a reproducible framework for embedding mechanical evaluation into experimental and translational oncology workflows.

Future developments will focus on expanding this strategy to additional tumor types, incorporating dynamic cues (*e.g.*, flow, immune components), and exploring integration with diagnostic imaging and patient-specific models.

## Supporting information

Main text

## References

1. Arnold M, Morgan E, Rumgay H, et al. Current and future burden of breast cancer: Global statistics for 2020 and 2040. Breast. 2022 Dec;66:15–23.

2. Valastyan S, Weinberg Robert A. Tumor Metastasis: Molecular Insights and Evolving Paradigms. Cell. 2011 2011/10/14/;147(2):275–292.

3. Massey A, Stewart J, Smith C, et al. Mechanical properties of human tumour tissues and their implications for cancer development. Nature Reviews Physics. 2024 2024/04/01;6(4):269–282.

4. Broders-Bondon F, Nguyen Ho-Bouldoires TH, Fernandez-Sanchez M-E, et al. Mechanotransduction in tumor progression: The dark side of the force. Journal of Cell Biology. 2018;217(5):1571–1587.

5. Peng H, Chao Z, Wang Z, et al. Biomechanics in the tumor microenvironment: from biological functions to potential clinical applications. Experimental Hematology & Oncology. 2025 2025/01/11;14(1):4.

6. Bao L, Kong H, Ja Y, et al. The relationship between cancer and biomechanics. Front Oncol. 2023;13:1273154.

7. Mierke CT. Viscoelasticity Acts as a Marker for Tumor Extracellular Matrix Characteristics. Front Cell Dev Biol. 2021;9:785138.

8. Radman BA, Alhameed AMM, Shu G, et al. Cellular elasticity in cancer: a review of altered biomechanical features. J Mater Chem B. 2024 Jun 5;12(22):5299–5324.

9. Nuckhir M, Withey D, Cabral S, et al. State of the Art Modelling of the Breast Cancer Metastatic Microenvironment: Where Are We? J Mammary Gland Biol Neoplasia. 2024 Jul 16;29(1):14.

10. Sixt M, Lämmermann T. In vitro analysis of chemotactic leukocyte migration in 3D environments. Cell Migration: Springer; 2011. p. 149–165.

11. Lämmermann T, Bader BL, Monkley SJ, et al. Rapid leukocyte migration by integrin-independent flowing and squeezing. Nature. 2008;453(7191):51–55.

12. Friedl P, Bröcker EB. The biology of cell locomotion within three-dimensional extracellular matrix. Cellular and molecular life sciences CMLS. 2000;57(1):41–64.

13. Sant S, Johnston PA. The production of 3D tumor spheroids for cancer drug discovery. Drug Discovery Today: Technologies. 2017;23:27–36.

14. Ng KW, Leong DTW, Hutmacher DW. The challenge to measure cell proliferation in two and three dimensions. Tissue engineering. 2005;11(1-2):182–191.

15. Rhodes NP, Srivastava JK, Smith RF, et al. Metabolic and histological analysis of mesenchymal stem cells grown in 3-D hyaluronan-based scaffolds. Journal of Materials Science: Materials in Medicine. 2004;15(4):391–395.

16. Banche-Niclot F, Lim J, McCulloch P, et al. Mesenchymal Stromal Cell Immunomodulatory Potential for Orthopedic Applications can be fine-tuned via 3D nano-engineered Scaffolds. Current Stem Cell Reports. 2024 2024/12/01;10(4):65–76.

17. Chuprin J, Buettner H, Seedhom MO, et al. Humanized mouse models for immuno-oncology research. Nat Rev Clin Oncol. 2023 Mar;20(3):192–206.

18. Li W, Zhou Z, Zhou X, et al. 3D Biomimetic Models to Reconstitute Tumor Microenvironment In Vitro: Spheroids, Organoids, and Tumor-on-a-Chip. Adv Healthc Mater. 2023 Jul;12(18):e2202609.

19. Muthuswamy SK, Brugge JS. Organoid Cultures for the Study of Mammary Biology and Breast Cancer: The Promise and Challenges. Cold Spring Harb Perspect Med. 2024 Jul 1;14(7).

20. Silva J, Oliveira PA, Duarte JA, et al. Mammary Cancer Models: An Overview from the Past to the Future. In Vivo. 2025 Jan-Feb;39(1):1–16.

21. Andolfi L, Greco SLM, Tierno D, et al. Planar AFM macro-probes to study the biomechanical properties of large cells and 3D cell spheroids. Acta Biomaterialia. 2019;94:505–513.

22. Sakuma S, Sato A, Kojima N, et al. Force sensor probe using quartz crystal resonator with wide measurement range for mechanical characterization of HepG2 spheroid. Sensors and Actuators A: Physical. 2017;265:202–210.

23. Liu H, Wen J, Xiao Y, et al. In situ mechanical characterization of the cell nucleus by atomic force microscopy. ACS nano. 2014;8(4):3821–3828.

24. Stewart DC, Rubiano A, Dyson K, et al. Mechanical characterization of human brain tumors from patients and comparison to potential surgical phantoms. PloS one. 2017;12(6):e0177561.

25. Jaiswal D, Cowley N, Bian Z, et al. Stiffness analysis of 3D spheroids using microtweezers. PloS one. 2017;12(11):e0188346.

26. Ferraro R, Caserta S, Guido S. A Low-Cost, User-Friendly Rheo-Optical Compression Assay to Measure Mechanical Properties of Cell Spheroids in Standard Cell Culture Plates. Advanced Materials Technologies. 2024 2024/01/05;n/a(n/a):2301890.

27. Ferraro R, Guido S, Caserta S, et al. i-Rheo-optical assay: Measuring the viscoelastic properties of multicellular spheroids. Materials Today Bio. 2024;26:101066.

28. Özkaya N, Leger D, Goldsheyder D, et al. Mechanical Properties of Biological Tissues. In: Özkaya N, Leger D, Goldsheyder D, et al., editors. Fundamentals of Biomechanics: Equilibrium, Motion, and Deformation. Cham: Springer International Publishing; 2017. p. 361–387.

29. Metzger W, Sossong D, Bächle A, et al. The liquid overlay technique is the key to formation of co-culture spheroids consisting of primary osteoblasts, fibroblasts and endothelial cells. Cytotherapy. 2011 Sep;13(8):1000–12.

30. Samani A, Plewes D. An inverse problem solution for measuring the elastic modulus of intact ex vivo breast tissue tumours. Phys Med Biol. 2007 Mar 7;52(5):1247–60.

31. Griffin M, Premakumar Y, Seifalian A, et al. Biomechanical Characterization of Human Soft Tissues Using Indentation and Tensile Testing. J Vis Exp. 2016 Dec 13(118).

32. Sutherland RM. Cell and environment interactions in tumor microregions: the multicell spheroid model. Science. 1988 Apr 8;240(4849):177–84.

33. Choi SH, Kim YH, Quinti L, et al. 3D culture models of Alzheimer’s disease: a road map to a “cure-in-a-dish“. Mol Neurodegener. 2016 Dec 9;11(1):75.

34. Yamada KM, Cukierman E. Modeling tissue morphogenesis and cancer in 3D. Cell. 2007 Aug 24;130(4):601–10.

35. Isvilanonda V, Li EY, Williams ED, et al. Subject-specific material properties of the heel pad: An inverse finite element analysis. Journal of Biomechanics. 2024 2024/03/01/;165:112016.

36. Gatt R, Vella Wood M, Gatt A, et al. Negative Poisson’s ratios in tendons: An unexpected mechanical response. Acta Biomater. 2015 Sep;24:201–8.

37. Islam MT, Tang S, Liverani C, et al. Non-invasive imaging of Young’s modulus and Poisson’s ratio in cancers in vivo. Scientific Reports. 2020 2020/04/29;10(1):7266.

38. Zanoni M, Piccinini F, Arienti C, et al. 3D tumor spheroid models for in vitro therapeutic screening: a systematic approach to enhance the biological relevance of data obtained. Scientific Reports. 2016 2016/01/11;6(1):19103.

39. Agrawal A, Lasli S, Javanmardi Y, et al. Stromal cells regulate mechanics of tumour spheroid. Materials Today Bio. 2023 2023/12/01/;23:100821.

40. Aung A, Davey SK, Theprungsirikul J, et al. Deciphering the Mechanics of Cancer Spheroid Growth in 3D Environments through Microfluidics Driven Mechanical Actuation. Adv Healthc Mater. 2023 Jun;12(14):e2201842.

41. Stylianopoulos T, Martin JD, Snuderl M, et al. Coevolution of solid stress and interstitial fluid pressure in tumors during progression: implications for vascular collapse. Cancer Res. 2013 Jul 1;73(13):3833–41.

42. Kontomaris SV, Stylianou A, Nikita KS, et al. Determination of the linear elastic regime in AFM nanoindentation experiments on cells [Article]. Materials Research Express. 2019 Nov;6(11):11.

43. Ciccone G, Dobre O, Gibson GM, et al. what caging force cells feel in 3D hydrogels: A rheological perspective. Advanced healthcare materials. 2020;9(17):2000517.

44. Ferraro R, Guido S, Caserta S, et al. Compressional stress stiffening & softening of soft hydrogels - how to avoid artefacts in their rheological characterisation. Soft Matter. 2023.

45. Elosegui-Artola A. The extracellular matrix viscoelasticity as a regulator of cell and tissue dynamics. Current opinion in cell biology. 2021;72:10–18.

