## Supplementary material for "Mechanical Fingerprints in Breast Cancer Research: A Multimodal Experimental Approach": Main text

### Supplementary information

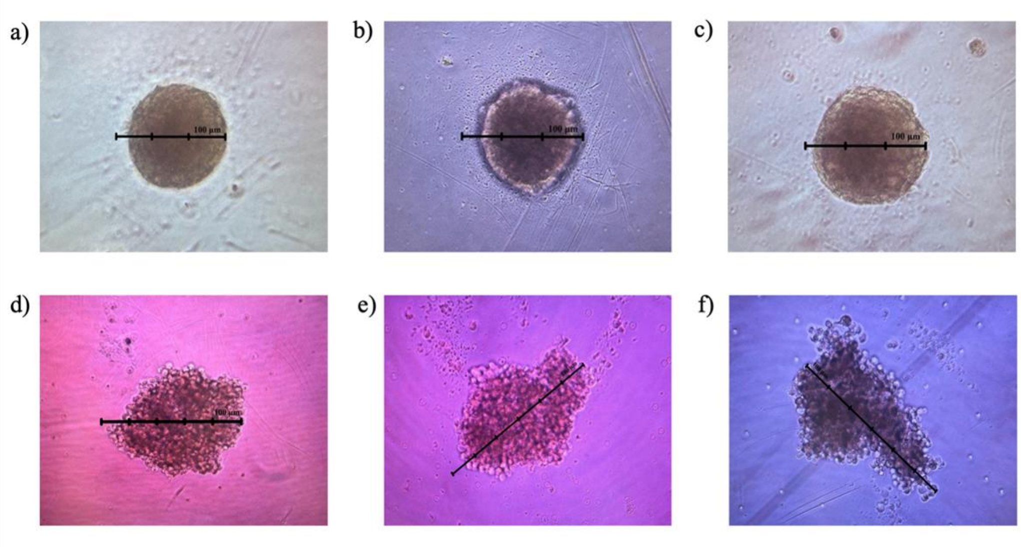


***Figure S1***. *Representative bright-field images of FP-MSC (**a-c**) and MDA-MB-231-GFP (**d-f**) spheroids used for size evaluation. Scale bar: 100 µm.*
